# Agentic systems are adept at solving well-scoped, verifiable problems in computational biology

**DOI:** 10.64898/2026.04.06.716850

**Authors:** Surag Nair, Laura Gunsalus, Brian Orcutt-Jahns, Jordan Rossen, Avantika Lal, Carlo De Donno, Muhammed Hasan Çelik, Kipper Fletez-Brant, Xiaoman Xie, Hector Corrada Bravo, Gokcen Eraslan

**Affiliations:** Computational Sciences Center of Excellence, Genentech, South San Francisco, CA, USA; Computational Sciences Center of Excellence, F. Hoffmann-La Roche, Basel, Switzerland

## Abstract

We introduce CompBioBench, a benchmark of 100 diverse tasks for evaluating agentic systems in computational biology. Unlike mathematics and programming, which more readily admit systematic verification, biological data are inherently noisy and open to interpretation. To enable objective evaluation without reducing tasks to prescriptive checklists, we propose a new benchmark construction strategy based on synthetic/augmented data and metadata scrambling/scrubbing of real datasets to create challenging problems that have a single ground-truth answer and require multi-step reasoning, tool use, bespoke code, and interaction with real-world external resources. The benchmark spans genomics, transcriptomics, epigenomics, single-cell analysis, human genetics, and machine learning workflows. Questions are curated by domain experts to cover a broad range of skills with varying difficulty. We evaluate leading general-purpose agentic systems starting from a bare-minimum environment, requiring them to fetch data and tools as needed to solve each problem. We find strong end-to-end performance, with Codex CLI (GPT 5.4) reaching 83% accuracy, Gemini CLI (3.1 Pro) reaching 82%, Claude Code (Opus 4.6) reaching 81%, and Claude Code (Opus 4.7) reaching 78%. On the hardest questions, Claude Code (Opus 4.6) reaches 69%, Codex CLI (GPT 5.4) reaches 59%, and Gemini CLI (3.1 Pro) reaches 49%. CompBioBench provides a practical testbed for measuring the progress of agentic systems in computational biology and for guiding future benchmark design. Data and a public leaderboard are available at https://huggingface.co/collections/Genentech/compbiobench-v1.

## Introduction

Computational biology is a highly interdisciplinary field, drawing on computer science, statistics, machine learning, genetics, and molecular biology^1,2^. It spans a wide range of problems and tasks, including exploratory data analysis, processing and interpreting high-throughput sequencing assays (e.g., RNA-seq, ATAC-seq, and single-cell modalities), variant analysis and interpretation, metagenomics, comparative genomics, proteomics, and integrative multi-omics. Practitioners in this field routinely navigate heterogeneous file formats, write and debug scripts and pipelines, and rely on a long tail of specialized tools, databases, and web resources.

Recent general-purpose agentic systems like Anthropic’s Claude Code and OpenAI’s Codex CLI have demonstrated the ability to tackle complex, open-ended tasks that combine reasoning and planning with tool use, web search, and computational environment setup^3,4^. More broadly, recent general-purpose systems have achieved strong results on demanding mathematics and programming tasks^5,6^. Benchmarking such agents in computational biology is challenging because many biologically meaningful analyses are noisy, underdetermined, and do not admit a single unambiguous ground-truth answer. Prior efforts to evaluate agents on real-world scientific and bioinformatics workflows such as BixBench^7^, Biomni-Eval1^8^, BioAgent Bench^9^, and scBench^10^ provide important early signals. However, these benchmarks focus on narrow slices of the domain, include multiple closely related tasks or underspecified questions, or specify the intended approach too explicitly, which reduces the need for autonomous strategy selection.

Here, we introduce CompBioBench, a benchmark consisting of 100 diverse computational biology questions with a focus on genomics, transcriptomics, epigenomics, single-cell analysis, human genetics, and machine learning applications. CompBioBench is designed to address these limitations: it spans multiple subfields and tests each concept in at most 2–3 questions to maximize diversity. To enable objective evaluation without reducing tasks to prescriptive checklists, we rely on data augmentation and synthesis, as well as scrambling or scrubbing metadata of real datasets, to construct questions that require multi-step reasoning, tool use, bespoke code, and interfacing with real-world external resources. CompBioBench includes tasks as diverse as identifying contaminants in RNA-seq, quantifying enzyme bias in ATAC-seq, inferring overexpressed TFs from chromatin accessibility data, aligning Perturb-seq datasets, detecting sample swaps in understudied species, demultiplexing scRNA-seq, inferring ancestry from personal variants, and designing mRNA sequences to optimize properties predicted by published machine learning models.

We benchmark Claude Code, Gemini CLI, and Codex CLI with different underlying models on CompBioBench. These general-purpose agents generally exhibited an impressive ability to solve complex, multi-step problems, with the best agent achieving 83.3% accuracy. We highlight a few selected examples, summarize lessons learned from designing this benchmark, and provide guidelines for future benchmarks for agentic systems in computational biology.

### Benchmark design/philosophy

CompBioBench aims to cover a diverse set of tasks encountered in computational biology (Methods). Although ambiguity is pervasive in biological data analysis, we designed questions with a single, unambiguous answer. At the same time, we avoided being prescriptive about the exact tools, workflows, or parameters required to reach that answer, and we limited repeated testing of any single concept or tool to no more than 2–3 problems. Because real biological data are inherently noisy and open to interpretation, establishing clear ground truth is often difficult. To enable objective evaluation without making tasks trivial, we made extensive use of synthetic and augmented data, as well as metadata recovery tasks (e.g., by scrambling or scrubbing metadata) on real data.

As an example of synthetic/augmented data, in question `contaminated-rna-q1`, we mixed human RNA-seq reads with RNA-seq reads from a distant species in specific proportions. Reads were trimmed to the same length, shuffled, and stripped of identifying FASTQ header information. The agent was then tasked with identifying the contaminating species. As an example of metadata recovery, in `sample-swap-atac-q1` we began with a bulk ATAC-seq counts matrix of axolotl at 10 kb resolution^11^, and swapped two closely related tissue labels. The agent was asked to determine whether a label-swapping event occurred and, if so, to return the names of the two swapped cell types. While such questions can be straightforward to construct, solving them often requires multi-step reasoning, reference data downloads/lookups, tool-calling, and bespoke code. By intentionally omitting the analytical strategy, these tasks test an agent’s algorithmic independence and its ability to navigate less common workflows. We complemented these with questions involving more routine analyses and data-retrieval tasks.

A crucial design decision concerned expectations around the base environment, tool setup, and data availability. For each question, the agents are provided with a bare-minimum compute environment and no data beyond the input files, with the expectation that they fetch required resources from the internet and modify the environment as needed. We believe this better reflects day-to-day workflows, which often involve a long tail of niche tools with small user bases and tools that are not actively maintained. To explicitly evaluate this capability, we included some “Tooling” questions that test an agent’s ability to initialize non-trivial environments. For example, the `encode-atac-pipeline-q1` question requires the agent to fetch code for the ENCODE ATAC-seq processing pipeline (v2.2.3)^12^, set up a compute environment, determine how to instantiate the pipeline, run it on provided inputs, navigate the output directory, and report the number of peaks called at the end of the deterministic pipeline.

Evaluation is binary, based on an exact string match between the agent’s answer and the ground truth. Expected output formats (e.g., exact comma-separated strings or specific casing) are clearly specified in each question. Numerical questions are designed to allow limited tolerance (e.g., by rounding to the nearest integer). In a few questions, multiple-choice options are explicitly provided.

CompBioBench contains a total of 100 questions with a focus on computational biology, spanning genomics, epigenomics, transcriptomics, and single-cell biology, along with a smaller set covering human genetics and machine learning workflows (Figure 1A, Supplementary Table 1). Nearly half of the questions use synthetic/augmented data or metadata recovery, while the remainder focus on routine analyses, data retrieval, and tooling (Figure 1B). Collectively, the benchmark targets a wide range of skills (Supplementary Figure 1). Furthermore, the questions were designed to have varying levels of difficulty in terms of logical reasoning and steps required, as reflected in the contributor-annotated difficulty level scores on a 1–5 scale (Figure 1C).

**Figure 1.**
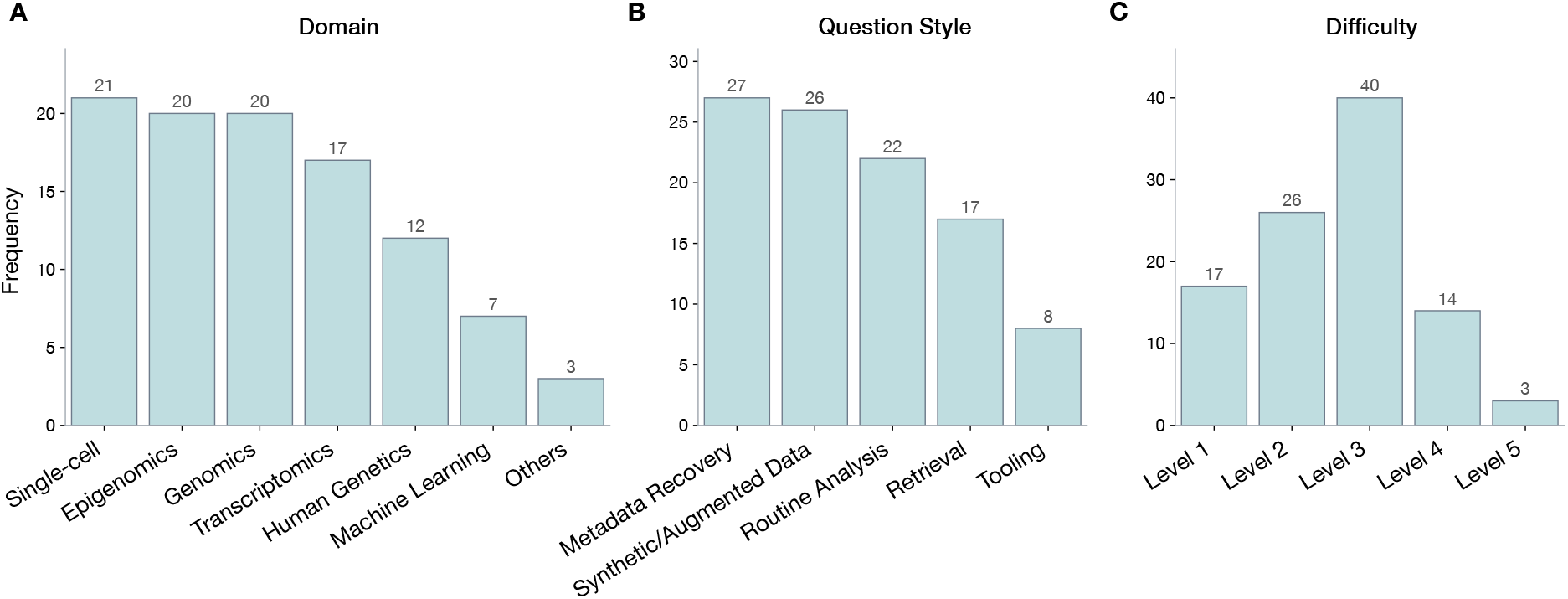
Split of questions in CompBioBench by **A)** domain, **B)** question style, and **C)** contributor-annotated difficulty level of problems. Each question is assigned exactly one domain and one question style.

Additional case studies illustrating the scope and complexity of these tasks are provided in Selected Examples and Supplementary Selected Examples.

### Performance of agentic systems

As a baseline, we ran two non-agentic, single-pass LLMs through API calls (Methods). We did not provide any input files in this run. As expected, these runs yielded poor performance: Claude Opus 4.6 achieved 3.7% accuracy while ChatGPT 5.2 achieved 5.3% (Figure 2A). The non-zero accuracy reflects random guessing on questions with a limited set of multiple-choice options, which confirms that the benchmark cannot be solved through memorized knowledge alone.

**Figure 2.**
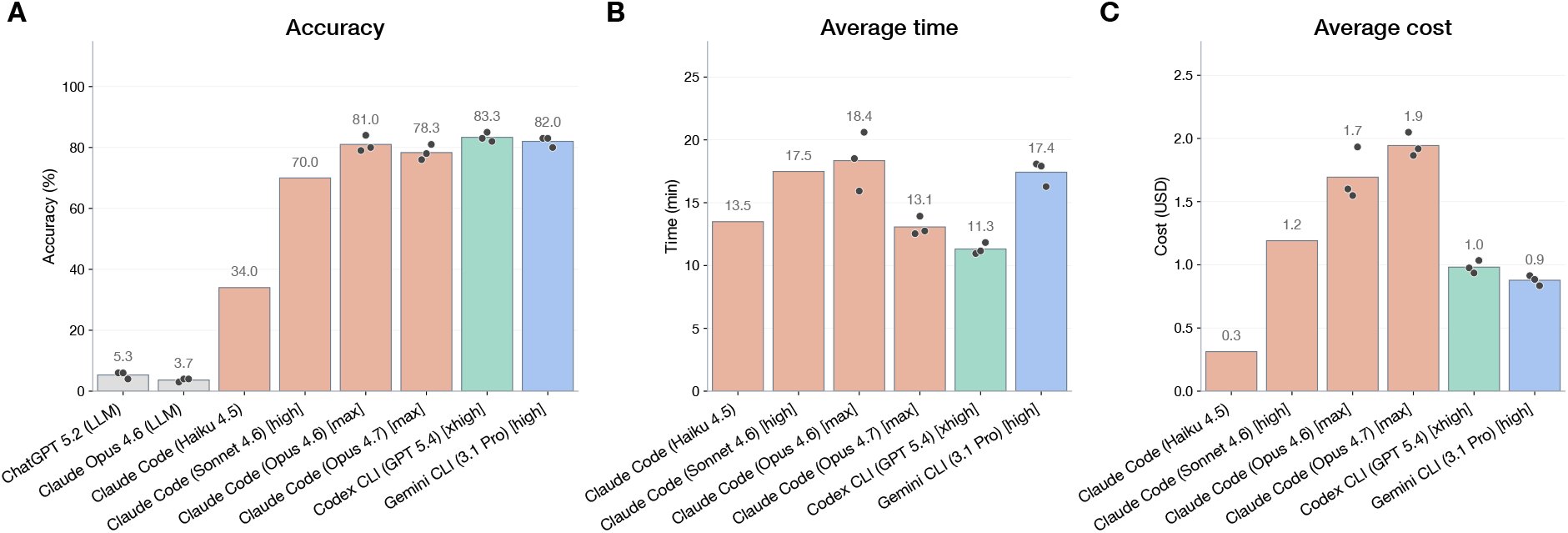
**A)** Average accuracy of LLM-only models and agentic systems across all questions, **B)** Wall-clock time taken per question, **C)** Estimated cost per question. Dots represent values for individual runs. The labels in square brackets indicate the reasoning level of the agents, which was set to the highest level possible. Bars for agentic systems are colored by provider.

We evaluated general-purpose agentic systems using a common runner script that constructs a standardized prompt, executes each agent CLI in an isolated per-question workspace and environment, and saves execution traces, token usage, cost, and other details (Methods). For each question, the initial environment consisted of the bare-minimum scientific and computational biology tools. The agent was given only the corresponding input files within this workspace, and no resource files (e.g., genome files, alignment indices). Web access was enabled and explicitly encouraged. Agents were also encouraged to download files, install tools, and update their environment as needed. When multiple reasoning levels were available for an agent, we selected the highest available setting.

Among the leading general-purpose agentic systems, Codex CLI (GPT 5.4) achieved the highest average accuracy at 83.3%, Gemini CLI (3.1 Pro) achieved 82.0%, and Claude Code (Opus 4.6) scored 81.0% (Figure 2A). Claude Code (Opus 4.7) scored 78.3%, underperforming Claude Code (Opus 4.6) despite being a newer model. We observed substantial variability: across the three runs of Codex CLI (GPT 5.4), 73% of questions were answered correctly on all three attempts, and 92% were answered correctly at least once. For Claude Code (Opus 4.6), 73% of questions were answered correctly on all three attempts, and 86% were answered correctly at least once. Performance degraded for smaller models. Claude Code (Sonnet 4.6) scored 70.0%, and Claude Code (Haiku 4.5) scored 34.0%. Among the leading agents, Claude Code (Opus 4.6) was the slowest, taking approximately 18 minutes per question on average, while Codex CLI (GPT 5.4) was the fastest at 11 minutes (Figure 2B). Claude Code (Opus 4.7) was the most expensive in our setup, consuming an average of USD 1.9 per question (Figure 2C). Gemini CLI (3.1 Pro) was the cheapest among the leading agents at USD 0.9 per question. We focus subsequent analysis on the three top-performing agents.

Performance differed across domains and difficulty levels (Supplementary Figure 2A,B). Codex CLI (GPT 5.4) scored 97% on 20 Genomics questions and achieved 70%+ accuracy in the remaining domains. All agents showed clear performance degradation as contributor-annotated difficulty increased. Codex CLI (GPT 5.4) achieved 100% accuracy on Level 1 questions, compared with 98% for Gemini CLI (3.1 Pro), and 90% for Claude Code (Opus 4.6). On the hardest questions (17 questions from Levels 4 and 5), Claude Code (Opus 4.6) scored 69%, outperforming Codex CLI (GPT 5.4) at 59% and Gemini CLI (3.1 Pro) at 49%. For all agents, time and cost increased with contributor-annotated difficulty level (Supplementary Figure 2C,D). For example, Codex CLI (GPT 5.4) took an average of 3 minutes, 6.5 minutes, 12.5 minutes, and 25 minutes for Level 1, 2, 3, and 4/5 questions, respectively.

We inspected agent traces qualitatively. Agents typically began by inspecting inputs and attempting an initial analysis, and often pivoted to alternate approaches when the first attempt failed. We observed frequent and largely frictionless use of external resources (APIs, web pages, code repositories, and public data portals) and routine installation or configuration of specialized tools when needed. In some cases, agents re-implemented task-relevant logic directly in code rather than relying on an existing library. Occasionally, we observed that once they arrived at an answer, they attempted to verify it using alternate approaches. In general, we found their breadth of resource and tool knowledge to be fairly comprehensive, with agents independently selecting and installing specialized tools such as Sourmash^13^, Vireo^14^, WhatsHap^15^, CellTypist^16^, and Chromosight^17^. The agents also routinely monitored their processes, occasionally terminated running code, and restarted it after optimizing for speed and/or memory.

### Selected examples

In this section, we highlight selected examples of questions and summarize the approach taken by Codex CLI (GPT 5.4), Gemini CLI (3.1 Pro), and Claude Code (Opus 4.6) on those tasks. Additional examples are provided in Supplementary Selected Examples.

**`subtype-inflammation-q1`**

**Q**. subtype.inflammation.q1.1.mtx.gz and subtype.inflammation.q1.2.mtx.gz contain raw scRNA counts matrices from retinal samples from two eyes of an individual (one per file, each column is a unique barcode that passes QC). subtype.inflammation.q1.genes.txt.gz is the list of common genes. One of the cell types is inflamed in one eye compared to the other. What is this cell type? Return only the name of the inflamed cell type from this list: astrocyte, microglial cell, RPE cell, cone cell, rod cell, horizontal cell, bipolar cell, RGC, T cells, macrophage, Schwann cell, Pericyte, B cell, fibroblast, muller glia cell, endothelial cell.

**Inputs:** 2 unannotated RNA-seq counts matrices with filtered cells, and a list of genes.

**Domain:** Single-cell

**Question type:** Synthetic/augmented data

**Difficulty:** 3/5

In this question, we started from filtered, preprocessed scRNA-seq count matrices for both eyes of a single individual from Wang *et al*.^18^. We introduced a targeted alteration by augmenting counts for a set of genes corresponding to an inflammation-linked program, restricted to one specific cell type in one eye, and then removed all annotations.

Agents first installed Scanpy^19^ and performed a standard scRNA-seq workflow: normalization and feature selection, dimensionality reduction, and clustering. They defined marker genes for each of the cell types listed in the question and labeled the cell types. They defined a list of inflammation-associated genes. They then tested for within-cluster differential expression between eyes, identified the cell type with the strongest enrichment for the inflammation gene set, and sanity-checked the call by inspecting individual inflammation-program genes and known markers before returning the final cell type. Agents typically took between 6–15 minutes to solve this task. Claude Code (Opus 4.6), Gemini CLI (3.1 Pro), and Codex CLI (GPT 5.4) solved it correctly on all occasions.

**`read-proportions-q1`**

**Q**. read.proportions.q1.fa contains a synthetic genome with four chromosomes. read.proportions.q1.fq.gz contains 40,000 single-end 36 bp reads drawn from this genome. A read is generated by first picking a chromosome with probability pi =(pi_1, …, pi_4), then sampling a position uniformly along that chromosome, with some small error rate. Estimate pi and report four integers (percentages) in FASTA contig order (chr1-chr4), each rounded to the nearest 5; if needed, adjust the last value so the total is 100. Expected output format: 15,25,25,35.

**Inputs:** A genome FASTA with four chromosomes (3,000 bp each) and a read FASTQ file with reads sampled from the chromosomes.

**Domain:** Genomics

**Question type:** Synthetic/augmented data

**Difficulty:** 3/5

In this question, we constructed a synthetic genome with extensive homology between chromosomes and then sampled reads according to the generative process described above. This setup induces a substantial fraction of multi-mapping reads. The question is designed such that naive counting of primary alignments yields the wrong answer.

Agents showed mixed performance on this task, and individual agents varied across runs. In all cases, they first built an alignment index and mapped reads using default parameters. In failing runs, the agent did not account for multi-mappers and terminated early, reporting pi based on the distribution of primary alignments among mapped reads. Agents that solved the problem correctly identified the multi-mapping issue, implemented a bespoke Expectation–Maximization procedure to infer pi under ambiguous assignments, and returned the correct estimate. Codex CLI (GPT 5.4) solved the problem correctly on all attempts, taking between 7–9 minutes. Gemini CLI (3.1 Pro) also solved it correctly on all attempts, taking between 6–8 minutes. Claude Code (Opus 4.6) solved it correctly only once. The incorrect attempts ended in about 1 minute, whereas the correct attempt took 8 minutes. This suggests that agent performance on such subtle tasks can be brittle: failure often reflects not an inability to solve the problem, but premature stopping after a superficially plausible analysis.

**`overexpress-tf-q1`**

**Q**. We perturbed a primary cell culture with a cocktail of transcription factors (TFs). ATAC-seq is performed on the initial sample and 48 hours after perturbation. overexpress.tf.q1.ref.bed.gz and overexpress.tf.q1.perturb.bed.gz contain the filtered aligned fragment files (hg38) corresponding to the initial and perturbed samples, respectively. Exactly one of the TFs in the following list is part of the cocktail: FOXA1, ASCL1, TFAP2A, RUNX1, KLF4, PAX7, TCF7, REST, SPI1, GATA3. Identify which one.

**Inputs:** 2 ATAC-seq fragment files

**Domain:** Epigenomics

**Question type:** Metadata recovery

**Difficulty:** 3/5

This question uses a real ATAC-seq dataset of transcription factor (TF) overexpression^20^ and is conceptually straightforward: identify regions with increased accessibility after perturbation and infer the overexpressed TF via motif enrichment. Agents generally had little trouble identifying differentially accessible regions. Some installed MACS2^21^, called peaks in each condition, and then tested for differential peaks. Others avoided peak calling by aggregating accessibility in fixed-length windows across the genome.

The main failure mode arose at the motif enrichment step. The choice of background sequences strongly affected the result: runs that used non-differential peaks (or other unmatched accessible regions) as background returned the wrong TF because the differential peaks have a shifted GC composition. Claude Code (Opus 4.6) and Gemini CLI (3.1 Pro) solved this question correctly on all attempts, using HOMER for motif enrichment each time. These runs took between 35 and 60 minutes. Codex CLI (GPT 5.4) solved it correctly only once. In that run, it also used HOMER and finished in 13 minutes. In the other two attempts, it wrote a bespoke motif-enrichment pipeline that ultimately failed as it did not select a GC-matched set of background regions. This highlights a broader robustness issue for agents, as even straightforward genomics tasks can hinge on subtle but consequential analytical decisions.

**`saluki-setup-optimize-q1`**

**Q**. Set up the Saluki model detailed in https://doi.org/10.1186/s13059-022-02811-x and use it to optimize the 3’ UTR of the mRNA sequence GCCGCCACCATGGTGAGCAAGGGCGAGTAGTGTACATAATAAGGACT. Specifically, perform 3 rounds of directed evolution, trying every possible substitution in the 3’ UTR and choosing the best after each round. Always use the average of predictions across all human model folds. Make sure to use GPUs if available. Report the full sequence of the optimized mRNA.

**Inputs:** None

**Domain:** Machine Learning

**Question type:** Tooling

**Difficulty:** 4/5

This question tests the ability of agents to download and run a machine learning model from the wild. Although it is not especially reasoning-heavy, it poses substantial logistical challenges, including locating the correct repository, downloading associated data, navigating complex codebases and data dumps, and configuring the compute environment. Below, we highlight how Codex CLI (GPT 5.4) solved the task and the obstacles it encountered along the way.

To locate the model, the agent first performed a web search and discovered that Saluki^22^, a deep learning model that predicts mRNA half-life from raw sequence, was not distributed as a standalone GitHub repository, but instead integrated into `calico/basenji`. After cloning that repository, it searched the codebase for Saluki-related terms and uncovered a manuscript README pointing to a secondary repository. By scanning this secondary codebase for inference-related files, it isolated the exact script used in the paper for fold-averaged 3’ UTR predictions. To obtain the pretrained model weights, the agent parsed the raw markdown of the original README and extracted a Zenodo DOI embedded in an image badge. It then determined that the checkpoints were bundled within a massive 18.75 GB archive. Rather than downloading the full dataset, the agent used HTTP byte-range requests to selectively extract only the 50 required fold-ensemble models and parameter files, reducing the transfer payload to roughly 104 MB. Such optimizations would be non-trivial for most human users.

The agent installed TensorFlow, but encountered substantial hardware-visibility issues. The agent resolved this by programmatically locating the missing NVIDIA libraries within the Python site-packages directory and injecting them into the execution environment. To begin the optimization, the agent first segmented the RNA sequence into its 5’ UTR, CDS, and 3’ UTR. It then deployed a custom Python script to perform directed evolution on the input mRNA sequence. Claude Code (Opus 4.6) and Codex CLI (GPT 5.4) solved this question correctly on all attempts in 12–35 minutes. Gemini CLI (3.1 Pro) solved it correctly two out of three times. The failed run resulted from incorrect input encoding. The entire task would likely take 3–4 hours for human experts.

Additional examples are provided in Supplementary Selected Examples.

### Benchmark design lessons from CompBioBench

In this section, we summarize practical lessons from developing CompBioBench that may inform future benchmarks for agentic systems.

First, we found it difficult to simultaneously satisfy three constraints: (i) requiring multi-step reasoning and tool use, (ii) avoiding prescriptive solution paths, and (iii) maintaining an unambiguous ground-truth answer. Constructing a single high-quality question often required several iterations, and the most challenging questions were typically grounded in concrete datasets and failure modes contributors had encountered in their own work. We therefore expect that building broad, challenging benchmarks will benefit from collaboration across larger teams with complementary domain expertise.

Second, while we initially attempted to use LLMs to propose candidate tasks under these constraints, the resulting ideas were often narrow, repetitive, or too simplistic. In our experience, LLMs were more useful for rapid implementation and iteration once a targeted task concept existed (e.g., drafting scripts, suggesting improvements, and stress-testing edge cases). Improved prompting, especially prompts that refine partially specified but domain-grounded ideas under explicit constraints, may further improve LLM utility during question formation.

Third, a straightforward path to scaling benchmark coverage is to extend existing question types in CompBioBench using controlled variations in datasets, parameters, and model choices, while preserving clear evaluation criteria. Because we test each core concept in only a small number of related questions, systematic expansion can increase robustness and highlight distinct failure modes.

Finally, although CompBioBench focuses on well-scoped problems with defined answers, we believe similar synthetic/augmentation strategies can support more discovery-oriented evaluations^23,24^. Large datasets can be injected with hidden plausible signals (e.g., subtle expression programs in a subpopulation) as well as hidden implausible artifacts (e.g., label errors, swaps, or outliers). Given a high-level discovery-oriented prompt, agents can perform a thorough analysis of the dataset and provide a report. Using an LLM-as-judge approach, the reports can be evaluated on their ability to recover these seeded signals. This strategy will require care to ensure that injected signals or artifacts are sufficiently realistic and do not shift the data too far from the real distribution, which could otherwise make the task artificially easy or less representative of the intended discovery setting.

## Discussion

We present CompBioBench, a benchmark of 100 diverse computational biology tasks for evaluating agentic systems. Leading general-purpose agents perform strongly, with Codex CLI (GPT 5.4) reaching 83% accuracy, Gemini CLI (3.1 Pro) reaching 82%, Claude Code (Opus 4.6) achieving 81%, and Claude Code (Opus 4.7) achieving 78%. These systems demonstrate the ability to solve well-scoped but non-trivial problems end-to-end, including multi-step reasoning, data download and processing, and tool use. Performance degrades substantially for smaller models, with Claude Code (Sonnet 4.6) achieving 70% and Claude Code (Haiku 4.5) achieving 34%. On the hardest questions, Claude Code (Opus 4.6) reaches 69%, Codex CLI (GPT 5.4) reaches 59%, and Gemini CLI (3.1 Pro) reaches 49%.

Agents remain brittle on certain analyses. These failures often arise not from an inability to solve the problem in principle, but from getting sidetracked by plausible alternative approaches or stopping prematurely after an initial but incomplete analysis. On longer-horizon tasks, such errors can compound over time. In practice, reliable deployment will still benefit from decomposing analyses into reasonably sized chunks (for example, tasks requiring roughly 30–60 minutes of agent runtime), together with human oversight to audit intermediate outputs and redirect the analysis when needed. Some of the brittleness we observed may also be mitigated by increased inference-time compute at the expense of additional cost. For example, running multiple agents in parallel and having a meta-agent combine the results could improve consistency.

In our benchmark, we instantiated the agents with a bare-minimum compute environment, only relevant input files, and no instructions beyond the problem statement. However, alternate approaches include pre-installing a comprehensive range of tools and datasets^8^, or providing “skill” files to the agents that outline high-level analytical strategies^25,26^. An important future direction is to evaluate whether these alternatives add value in terms of performance, time savings, token usage, and overall cost.

Together, these results mark a step change in our understanding of the ability of general-purpose agentic systems to perform routine computational biology analyses. The pace of recent progress suggests that such general-purpose systems could become dependable analysts in the future. They are already highly capable at general scripting, data wrangling, tool discovery and installation, and process orchestration. They can identify optimizations beyond what most human experts would readily attempt. Benchmarking efforts like CompBioBench will help build trust in agentic systems by grounding evaluation in routine workflows and tasks that computational biologists are familiar with. We hope CompBioBench, and the synthetic/augmented data and metadata recovery paradigm it emphasizes, will support broader community-driven benchmark development.

### Limitations

The scope of questions tested in CompBioBench is limited by the expertise of the contributors. The benchmark does not exhaustively cover all subdomains and routine tasks in computational biology. By design, it does not cover tasks with substantial biological or methodological ambiguity, where multiple approaches may yield different yet valid answers. It tests the ability to use existing tools and write custom scripts. However, it does not test the use of user-supplied tools (e.g., via Model Context Protocol), the development of new data processing pipelines, or the discovery of novel algorithms. Evaluation focuses on the final answer and does not score partial credit or intermediate outputs/artifacts. Due to the time required per task (often several hours for difficult questions), we did not collect systematic human performance estimates. Many questions require retrieving external reference data or software. We have attempted to rely on durable, widely used resources (e.g., 1000 Genomes, ENCODE, GEO, and DOI-addressable artifacts) so that the benchmark remains useful over time, but some dependencies may still change or become unavailable.

## Data and code availability

We released all questions and associated files on Zenodo at https://doi.org/10.5281/zenodo.19443185. Code for running all agentic systems is available at https://github.com/Genentech/compbiobench-runner. A Hugging Face leaderboard to enable developers to evaluate their agents on CompBioBench is available at https://huggingface.co/spaces/Genentech/compbiobench-leaderboard-v1.

## Acknowledgements

We would like to thank Sara Mostafavi, Parisa Mazrooei, Camilo Espinosa Bernal, Omar Salem, Archit Verma, Hugues Van Assel, Kenneth Gao, Edward De Brouwer, Jan-Christian Hütter, Jin Liu, and David Garfield for their feedback and discussion.

## Supplementary figures

**Supplementary Figure 1.**
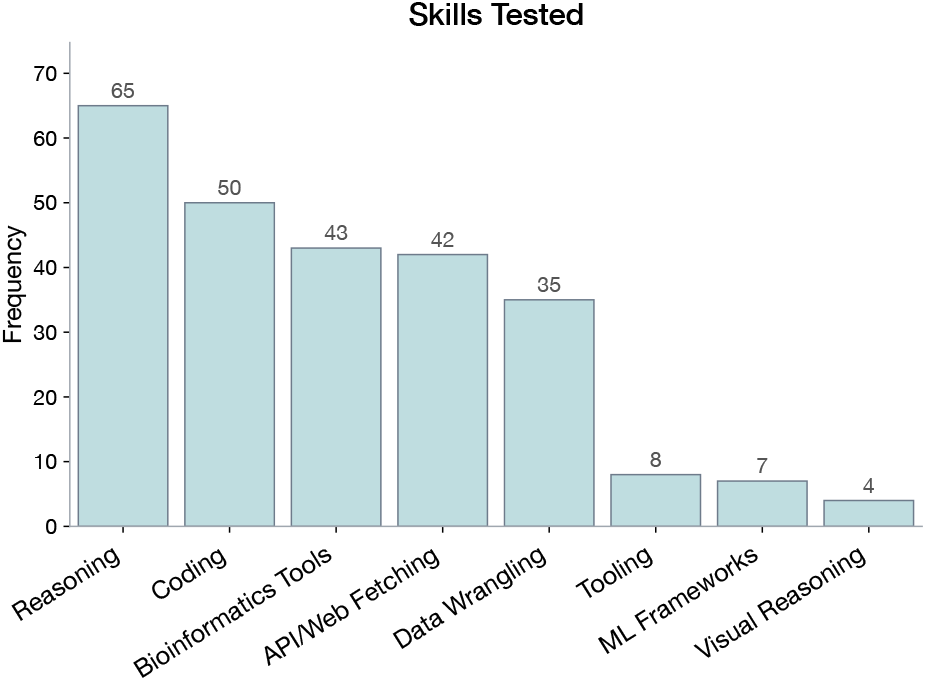
Split of questions by skills tested. Each question is assigned one or more skills.

**Supplementary Figure 2.**
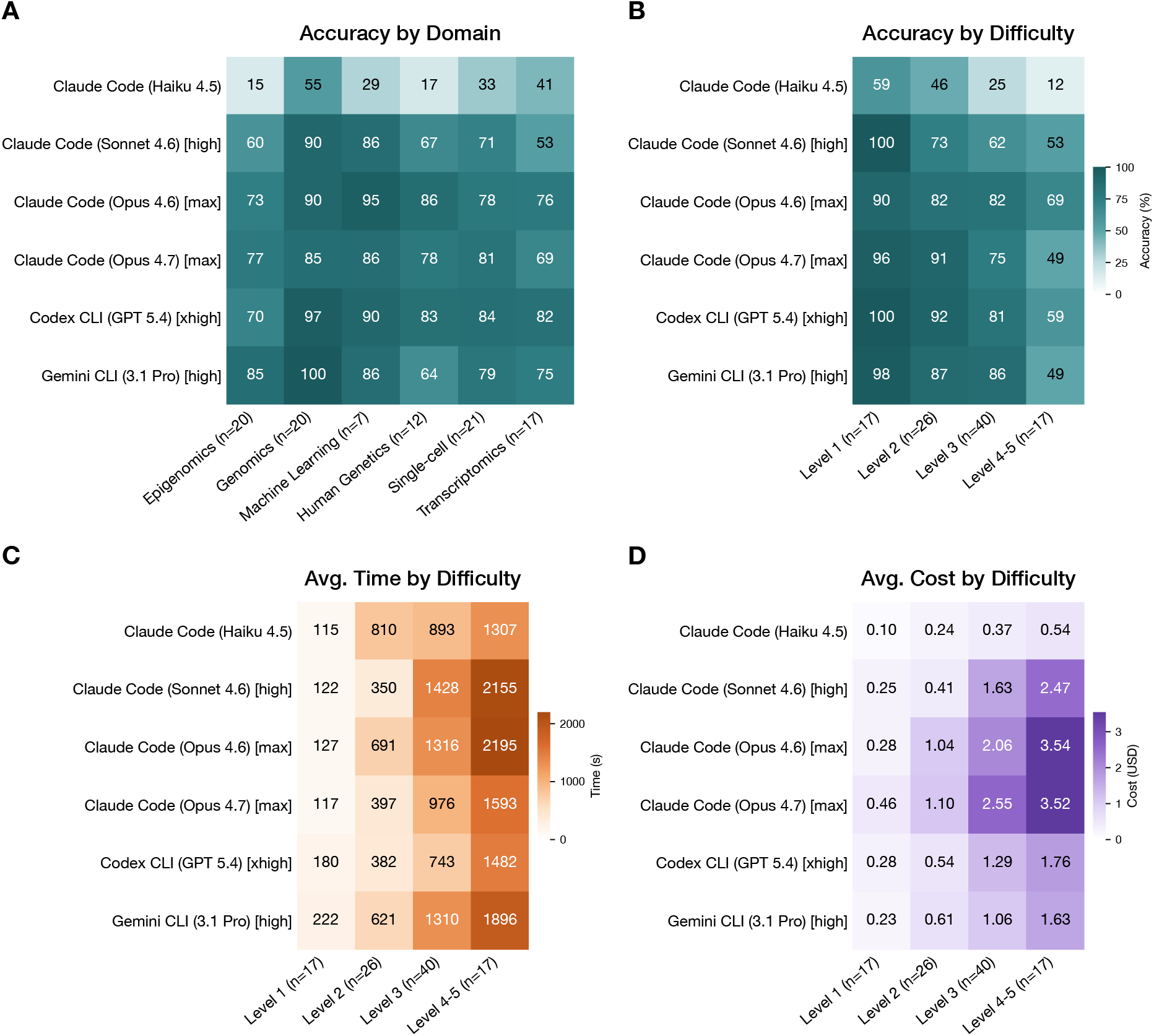
**A)** Accuracy of the agents stratified by domain. Domains with fewer than 5 questions were removed. **B)** Accuracy, **C)** average time taken, and **D)** average cost stratified by contributor-annotated difficulty. Difficulty Levels 4 and 5 were grouped together. Values are averaged over all runs for that agent.

## Supplementary selected examples

**`afgr-1000g-intersect-atac-q1`**

**Q**. From the 1000G Phase 3 (3202 individuals), find all individuals that also have ATAC-seq filtered BAMs in the African Functional Genomics Resource (AFGR). Using only publicly available data, collect the md5sums of the filtered BAMs for these individuals (GRCh38 aligned), write the md5sums (one per line) to a text file sorted alphabetically. Return the count of samples and the file’s md5sum as a single comma-separated line, e.g., 42,md5hash.

**Inputs:** None

**Domain:** Human Genetics

**Question type:** Retrieval

**Difficulty:** 4/5

This question tests the agents’ ability to navigate large public genomics databases, cross-reference sample identifiers, and programmatically filter complex metadata. As an example, Codex CLI (GPT 5.4) began by identifying that African Functional Genomics Resource^27^ data is hosted on the ENCODE portal. It used the ENCODE REST API, a resource it was familiar with. It downloaded the metadata TSV for all AFGR ATAC-seq experiments. It successfully filtered this dataset down to 100 GRCh38-aligned filtered BAM files by recognizing ENCODE’s specific file labeling distinctions (alignments vs. unfiltered alignments). Next, the agent needed the official list of 3202 individuals from the 1000 Genomes (1000G) Phase 3 project^28^. It directly queried the 1000G high-coverage FTP directories via curl, scraping the HTML index pages for keywords like “ped” and “sample” to pinpoint the exact demographic pedigree file.

It then resolved nomenclature discrepancies between the two datasets. While 1000G uses “NA” prefixes for Coriell cell line samples (e.g., NA19397), the ENCODE/AFGR metadata uses “GM” prefixes (e.g., GM19397). The agent recognized this alias pattern and mapped the IDs appropriately before performing the set intersection. It identified the overlapping individuals, extracted the corresponding md5sums from the metadata TSV, and generated the final answer. Agents typically took between 6–18 minutes to solve this task. Claude Code (Opus 4.6), Gemini CLI (3.1 Pro), and Codex CLI (GPT 5.4) solved it correctly on all occasions.

**`atac-doublet-q1`**

**Q**. atac.doublet.q1.bed.gz contains a fragment file from a scATAC-seq sample (hg38). Barcodes with low read counts have been filtered out. Look for doublets. Return the barcode of any one doublet, if any. Respond only with the label of the barcode in the format: “AAATGGAACGTTAAAG-1”, or “None”.

**Inputs:** a single ATAC-seq fragment file

**Domain:** Epigenomics

**Question type:** Synthetic/augmented data

**Difficulty:** 4/5

We started with a bulk cell-line ATAC-seq dataset and synthetically generated “cells” by assigning fragments to barcodes at random, so that each barcode has approximately the same number of fragments. We then introduced a single barcode exhibiting an excess of loci with >2 overlapping fragments. The agent is tasked with determining whether a doublet exists and, if so, returning the corresponding barcode.

The key insight is tied to the approach used by AMULET^29^. Unlike common scATAC-seq doublet detection methods that rely on co-localization of observed cells with simulated doublets, AMULET-style overlap-based signals can detect even homotypic doublets. No information about the underlying cell-type composition is provided, so the solution must rely on fragment-level patterns rather than expression- or cluster-based heuristics.

Codex CLI (GPT 5.4) and Claude Code (Opus 4.6) typically began by examining the per-barcode read-depth distribution, which failed to reveal outliers because fragment counts are intentionally balanced. After exploring a few other unsuccessful directions, successful runs pivoted to explicitly testing for abnormal fragment-overlap patterns. In these runs, the agents directly referenced AMULET, then either installed the package or reimplemented its core idea from scratch, allowing them to solve the question. Claude Code (Opus 4.6) solved it correctly on all attempts in 3–6 minutes. Codex CLI (GPT 5.4) solved it correctly once, in 5 minutes. In its incorrect attempts, it relied exclusively on synthetic doublet simulations across multiple genomic bin sizes, taking 20–30 minutes without finding the correct answer. Gemini CLI (3.1 Pro) solved it correctly on all three attempts under 4 minutes, and notably went straight to the AMULET-style overlap analysis without first detouring through read-depth checks or synthetic-doublet simulations. This example is notable since AMULET is not a standard part of single-cell ATAC-seq workflows. Furthermore, it highlights that agent success can depend critically on whether the right analytical hypothesis is considered early, as the agent can spend substantial effort pursuing plausible yet ultimately uninformative strategies.

**`hic-differential-loop-q1`**

**Q**. Consider the MicroC data for H1 and HFF cells from the following paper: https://doi.org/10.1016/j.molcel.2020.03.003. In the sub-compartment containing the NANOG gene (chr12:7629950-7809597), there is a differential loop between the two samples. Report the position of the loop (on chr12) in the following format start;end, with start<end rounded to the nearest 20000 and no commas.

**Inputs:** None

**Domain:** Epigenomics

**Question type:** Retrieval

**Difficulty:** 3/5

The agent was tasked with identifying the coordinates of a differential chromatin loop on chromosome 12 within the NANOG sub-compartment, specifically comparing Micro-C data from H1 and HFF cell lines published in Krietenstein *et al*.^30^. A human expert can solve this task in 5–10 minutes by navigating to the 4DNucleome (4DN) web browser^31^, visualizing the preprocessed contact matrices for the two cell lines side-by-side, and visually extracting the coordinates of the differential loop.

However, the agents approached this problem with divergent, non-visual methodologies. Codex CLI (GPT 5.4) solved the task correctly on all three attempts, taking between 7 and 22 minutes. Its general strategy more closely mirrored the human workflow: it programmatically interacted with the 4DN web portal and the HiGlass visualization APIs^32^. By extracting the localized image tiles for the target genomic region, the agent efficiently pinpointed the differential loop coordinates.

Gemini CLI (3.1 Pro) solved the task correctly twice, in one case in under 5 minutes. Both successful runs used the same method: rather than visualizing matrices, they discovered that 4DN hosts precomputed loop-call BEDPE files for each experiment set, downloaded those, and identified the differential loop. The failed run never found these files and instead tried to re-derive the loop from raw HiGlass contact-matrix tiles using ad hoc pixel-difference heuristics, which could not reliably pinpoint the coordinate.

In contrast, Claude Code (Opus 4.6) solved it correctly twice, but its runs took almost an hour. In one run, it brute-forced the problem by downloading 24 GB of raw multi-resolution contact matrices to computationally re-call the loops from scratch. In the other, it streamed the relevant data remotely, but lost significant time fighting environment dependency issues and compiling C++ libraries. Ultimately, as current CLI agents cannot natively browse web pages visually like humans, the exact analytical approach an agent commits to dictates the complexity and computational overhead of its solution.

## Methods

### Benchmark design process

We sourced questions from seven contributors. Each benchmark item consists of (i) a free-text question that specifies the task and optionally references one or more input files, (ii) the input files, if any, and (iii) a single string-valued answer used for evaluation. Tasks are independent of one another, though some can test a different concept with the same inputs or test related concepts with different inputs.

We initially attempted to use LLMs to propose candidate tasks and used these ideas as the basis for some questions. However, we eventually found the ideas to be narrow, repetitive, or too simplistic. Ideas for subsequent questions originated from concrete datasets and problems encountered by contributors in their own work. We used LLMs for rapid implementation and iteration once a targeted task concept existed (e.g., drafting scripts, suggesting improvements, and stress-testing edge cases).

When questions required external data resources, we attempted to rely on durable, widely used sources (e.g., 1000 Genomes, ENCODE, GEO, and DOI-addressable artifacts). We recorded all files used, their checksums, and documented where each resource was downloaded from and how it was processed. For questions that rely on specific software or data versions, we explicitly specify those versions in the question text.

We began building the benchmark with simpler problems and ran them through agentic systems to better understand current capabilities and common failure modes, which informed the design of more challenging multi-step questions. In some cases, we introduced small complications intended to reflect realistic analysis pitfalls that we expected agentic systems to account for, such as deliberate off-by-one issues or mismatched gene orders between two AnnData objects that must be compared.

For synthetic or augmented-data questions (including metadata scrambling/scrubbing tasks), the ground truths were well-defined by design. For retrieval, routine analysis, and tooling questions, contributors first solved the task to establish the expected answer and to verify that the required resources and software could be obtained and executed as intended.

Most problems were refined in a loop that combined question development and agent trace review. After an initial problem proposal, we inspected traces of agentic systems to ensure that solutions could not exploit unintended shortcuts (e.g., information leakage through file headers or other metadata). If a question exhibited avoidable ambiguity that led to inconsistent interpretation, we added clarifying details to the prompt and corrected any discrepancies in the expected output format specification. Problems that were not solved correctly by any agentic system were reviewed carefully for correctness. In these cases, we also attempted to solve the task ourselves from first principles to validate that the question was solvable as stated and that the ground-truth answer was correct.

Finally, we annotated each question with domain, question style, and the skills tested. Annotations were produced by the lead author with LLM assistance. No inter-annotator agreement was measured. For each question, contributors provided a difficulty rating on a 1–5 scale. These ratings were not extensively calibrated across questions and should be interpreted as approximate.

### Running non-agentic baselines

We ran Claude Opus 4.6 and ChatGPT 5.2 models using their respective APIs with default parameters. We ran each question three times through the models. We did not provide any files as input. For each question, the full prompt for the model was:

~~~
 QUESTION: {question}
 REFERENCE FILES (NOT PROVIDED IN THIS RUN):
 {file_paths}
 CONTEXT:
 - This is a non-agentic benchmark run.
 - No local files, attachments, or workspace files are available.
 - If the question depends on missing files, give your best possible guess from
 prior knowledge.
 OUTPUT FORMAT REQUIREMENTS:
 When you have determined the answer, end your response with EXACTLY:
 FINAL ANSWER:
 <your answer here>
 IMPORTANT FORMAT RULES:
 1. The text ‘FINAL ANSWER:’ MUST be in all caps with a colon.
 2. Put the answer on the line(s) immediately after ‘FINAL ANSWER:’.
 3. The answer MUST follow any format specified in the question.
 4. This must be the last part of your response.
~~~

### Running agentic systems

We ran each agentic benchmark question through a common runner script that (i) constructs a standardized prompt, (ii) executes the chosen agent CLI inside an isolated, per-question conda clone, (iii) saves both a human-readable trace and a machine-readable JSON record, and (iv) saves metadata including token usage and cost. A single base conda environment (`compbio-benchmark`) is created from an `environment.yml` file (conda-forge/bioconda, Python 3.11 plus basic scientific and bioinformatics tooling, and Node.js for CLIs), then cloned per question to ensure reproducibility across parallel runs. For each question, required input files are copied into an isolated “workspace” directory, and the agent CLI is launched with that workspace as its current working directory.

Agents were instructed not to access files outside the workspace, but this is only a behavioral constraint (the runner does not enforce a hard filesystem sandbox). We analyzed traces to ensure that this constraint was observed. Web access was allowed in the configuration (Codex `network_access = true`), explicitly encouraged in the prompt, and the runner’s trace parser records web search/fetch events alongside shell commands and file reads/edits. To reduce interactive friction, the runner can launch agents in “auto-approve” modes (e.g., Claude `--dangerously-skip-permissions`; Codex `--dangerously-bypass-approvals-and-sandbox`; Gemini `--yolo`), effectively granting write actions without per-step confirmation.

Each question has a hard wall-clock timeout: the runner waits on the subprocess with a per-question limit and, on timeout, kills the entire process group to stop any spawned children. We initially ran all agents with a time limit of 120 minutes. For all agents except Claude Code (Haiku 4.5), we re-ran questions that timed out on the first attempt with a 240-minute time limit and a clean start (i.e., without resuming the previous run). Questions not solved within this time limit were marked incorrect.

Outputs were extracted from each provider’s native structured output channel. For Claude and Gemini, the runner parsed the JSONL event stream and took the last non-empty line of the final response. For Codex, the final response was written to an output file and then read. Occasionally, when agents did not adhere to the specified output format (most commonly Haiku and Sonnet) but nevertheless provided the correct answer, we manually marked them as correct.

In our experiments, we used Claude Code (v2.1.87) with the following models: Opus 4.6 (`claude-opus-4-6`), Sonnet 4.6 (`claude-sonnet-4-6`), and Haiku 4.5 (`claude-haiku-4-5-20251001`). For Sonnet and Opus, we used the 1M-context versions. When available, we set reasoning effort to the highest supported level. Specifically, we used the `max` reasoning level for Opus and `high` for Sonnet via the `--effort` flag. Effort was not supported for Haiku. We ran Claude Code with Opus 4.6 three times to estimate variability between runs. These runs were performed between March 23 and March 31, 2026. We used Claude Code (v2.1.119) with Opus 4.7 (`claude-opus-4-7`) with `max` reasoning, and ran the agent three times between April 20 and April 24, 2026.

We used Codex CLI (v0.115.0) with GPT 5.4 (`gpt-5.4`). We used the `xhigh` reasoning level via the `model_reasoning_effort` parameter. We ran Codex CLI (GPT 5.4) three times between March 23 and March 31, 2026.

We used Gemini CLI (v0.39.1) with Gemini 3.1 Pro Preview (`gemini-3.1-pro-preview`). We used `high` reasoning level by setting `“thinkingLevel”:”HIGH”` in the Gemini `settings.json` file. We ran the agent three times, between April 22 and April 25, 2026.

Experiments were run on an NVIDIA DGX A100 node with 2× AMD EPYC 7742 64-core CPUs (256 threads), 2 TB RAM, and 8× NVIDIA A100 80 GB GPUs. For all plots and summary metrics in the paper, we clipped the maximum time per question at 7,200 seconds and the maximum cost per question at USD 10 to reduce the effect of outliers.

For each question, the full prompt for the agent was:

~~~
 QUESTION: {question}
 FILES: {workspace_file_paths}
 Note: All files are located in your current working directory (workspace).
 INSTRUCTIONS:
 - You have {timeout_minutes} minutes to complete this task
 - Do not read/access any other files outside the workspace.
 - Get any files or tools you need from the internet.
 - You are free to modify the current conda environment as needed.
 - Keep all scripts and intermediate data in the workspace only.
 OUTPUT CONTRACT (IMPORTANT):
 - Your final response will be graded by exact string match.
 - Return EXACTLY ONE LINE containing **ONLY** the final answer in the format
   required by the question.
 - Do NOT include any explanation, reasoning, labels, prefixes, markdown, code
   fences, citations, or extra whitespace lines.
 - Any extra text before or after the answer is incorrect.
 - Before sending your final response, verify it is exactly one line and
   nothing else.
~~~

The YAML file used to instantiate the conda environment for each question is:

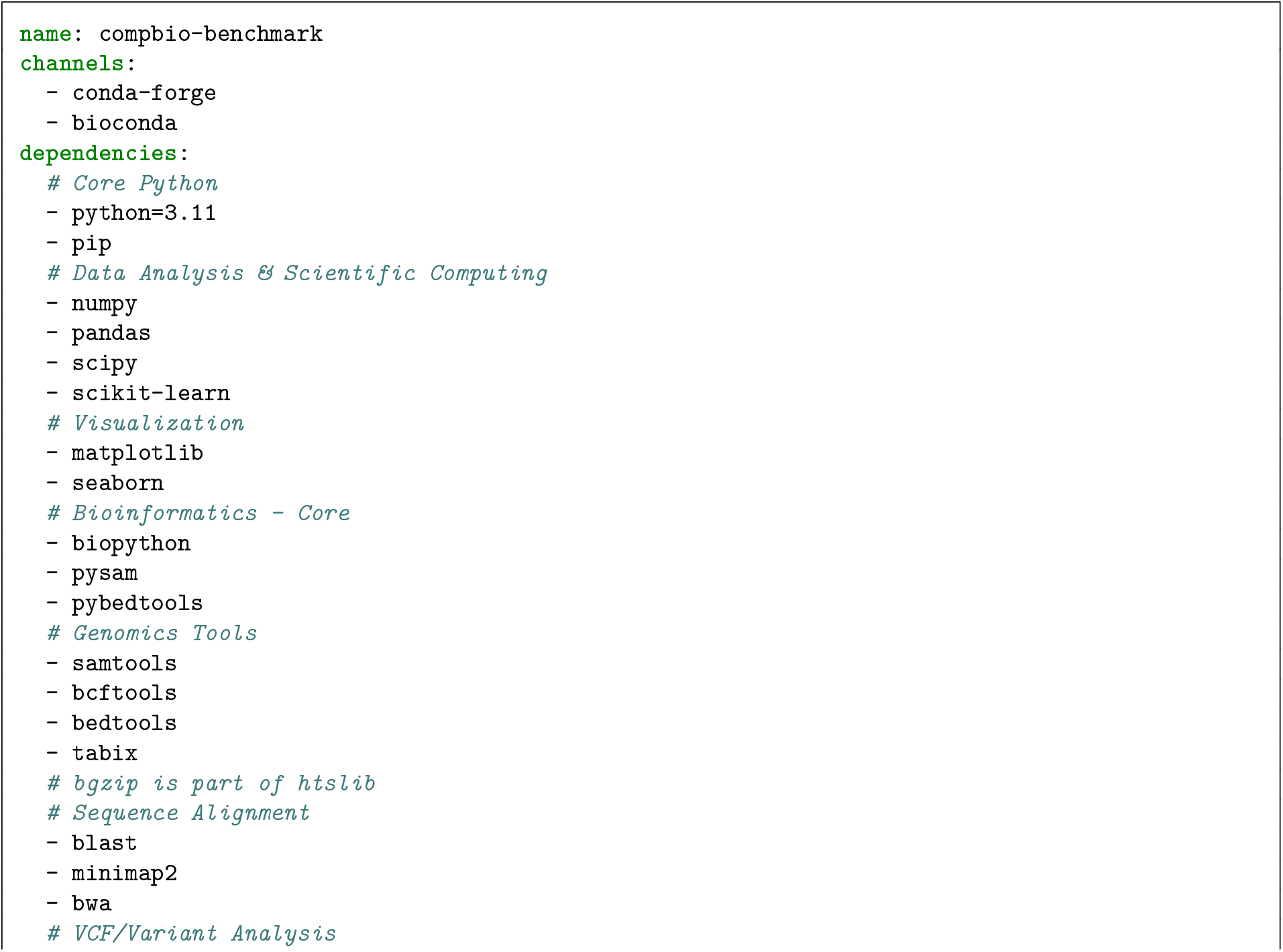
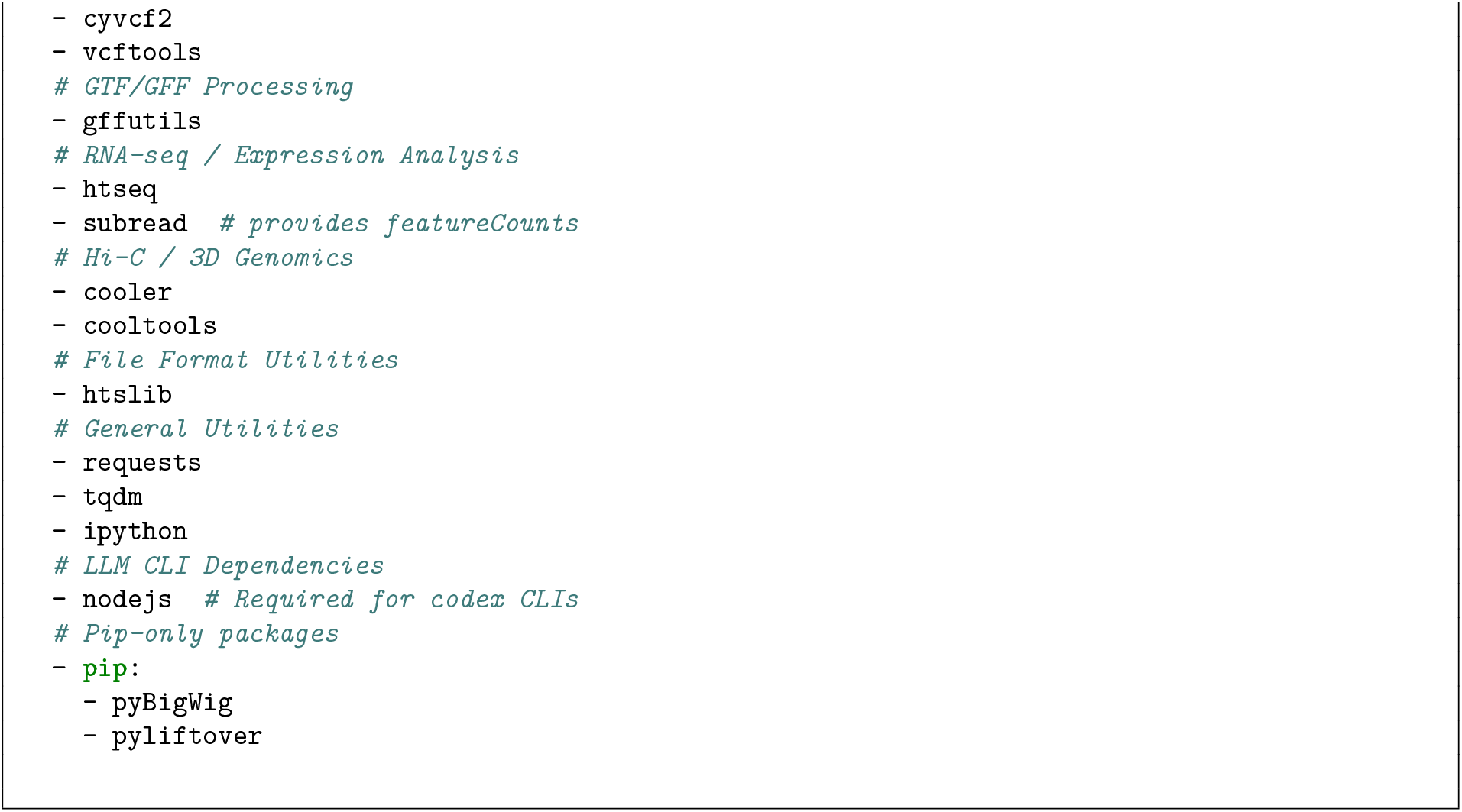

## Notes

https://doi.org/10.5281/zenodo.19443185

https://huggingface.co/spaces/Genentech/compbiobench-leaderboard-v1

https://github.com/Genentech/compbiobench-runner

### Competing Interest Statement

All authors are Genentech/Roche employees and may hold Roche shares or stock-based compensation; no other competing interests.

### Summary of Updates

Updated to include results of Claude Code (Opus 4.7) and Gemini CLI (3.1 Pro).

## References

1. Wendy Weijia Soon, Manoj Hariharan, and Michael P Snyder. High-throughput sequencing for biology and medicine. Molecular systems biology, 9:640, 2013.

2. Yiming Qin, Hari Krishna Yalamanchili, Jing Qin, Bin Yan, and Junwen Wang. The current status and challenges in computational analysis of genomic big data. Big data research, 2(1):12–18, 2015.

3. Mark Chen, Jerry Tworek, Heewoo Jun, Qiming Yuan, Henrique Ponde De Oliveira Pinto, Jared Kaplan, Harri Edwards, Yuri Burda, Nicholas Joseph, Greg Brockman, et al. Evaluating large language models trained on code. arXiv preprint arXiv:2107.03374, 2021.

4. Grégoire Mialon, Clémentine Fourrier, Thomas Wolf, Yann LeCun, and Thomas Scialom. GAIA: a benchmark for general AI assistants. In The Twelfth International Conference on Learning Representations, 2023.

5. Ahmed El-Kishky, Alexander Wei, Andre Saraiva, Borys Minaiev, Daniel Selsam, David Dohan, Francis Song, Hunter Lightman, Ignasi Clavera, Jakub Pachocki, et al. Competitive programming with large reasoning models. arXiv preprint arXiv:2502.06807, 2025.

6. Carlos E Jimenez, John Yang, Alexander Wettig, Shunyu Yao, Kexin Pei, Ofir Press, and Karthik Narasimhan. SWE-bench: Can language models resolve real-world GitHub issues? arXiv preprint arXiv:2310.06770, 2023.

7. Ludovico Mitchener, Jon M Laurent, Alex Andonian, Benjamin Tenmann, Siddharth Narayanan, Geemi P Wellawatte, Andrew White, Lorenzo Sani, and Samuel G Rodriques. BixBench: a comprehensive benchmark for LLM-based agents in computational biology. arXiv preprint arXiv:2503.00096, 2025.

8. Kexin Huang, Serena Zhang, Hanchen Wang, Yuanhao Qu, Yingzhou Lu, Yusuf Roohani, Ryan Li, Lin Qiu, Gavin Li, Junze Zhang, et al. Biomni: a general-purpose biomedical AI agent. bioRxiv, 2025.

9. Dionizije Fa, Marko Čuljak, Bruno Pandža, and Mateo Čupić. BioAgent Bench: An AI agent evaluation suite for bioinformatics. arXiv preprint arXiv:2601.21800, 2026.

10. Kenny Workman, Zhen Yang, Harihara Muralidharan, Aidan Abdulali, and Hannah Le. scBench: Evaluating AI agents on single-cell RNA-seq analysis. arXiv preprint arXiv:2602.09063, 2026.

11. Xiaoping Han, Hanyu Wu, Xueyi Wang, Daiyuan Liu, Yuting Fu, Lei Yang, Renying Wang, Peijing Zhang, Jingjing Wang, Lifeng Ma, et al. Modeling the vertebrate regulatory sequence landscape by UUATAC-seq and deep learning. Cell, 188(19):5343–5362, 2025.

12. Benjamin C Hitz, Lee Jin-Wook, Otto Jolanki, Meenakshi S Kagda, Keenan Graham, Paul Sud, Idan Gabdank, J Seth Strattan, Cricket A Sloan, Timothy Dreszer, et al. The ENCODE uniform analysis pipelines. Biorxiv, 2023.

13. N Tessa Pierce, Luiz Irber, Taylor Reiter, Phillip Brooks, and C Titus Brown. Large-scale sequence comparisons with sourmash. F1000Research, 8:1006, 2019.

14. Yuanhua Huang, Davis J McCarthy, and Oliver Stegle. Vireo: Bayesian demultiplexing of pooled single-cell RNA-seq data without genotype reference. Genome biology, 20(1):273, 2019.

15. Marcel Martin, Peter Ebert, and Tobias Marschall. Read-based phasing and analysis of phased variants with WhatsHap. In Haplotyping: Methods and protocols, pages 127–138. Springer, 2022.

16. César Domínguez Conde, Chao Xu, Louie B Jarvis, Daniel B Rainbow, Sara B Wells, Tamir Gomes, SK Howlett, O Suchanek, K Polanski, HW King, et al. Cross-tissue immune cell analysis reveals tissue-specific features in humans. Science, 376(6594):eabl5197, 2022.

17. Cyril Matthey-Doret, Lyam Baudry, Axel Breuer, Rémi Montagne, Nadège Guiglielmoni, Vittore Scolari, Etienne Jean, Arnaud Campeas, Philippe-Henri Chanut, Edgar Oriol, et al. Chromosight: a computer vision program for pattern detection in chromosome contact maps. bioRxiv, pages 2020–03, 2020.

18. Sean K Wang, Surag Nair, Rui Li, Katerina Kraft, Anusri Pampari, Aman Patel, Joyce B Kang, Christy Luong, Anshul Kundaje, and Howard Y Chang. Single-cell multiome of the human retina and deep learning nominate causal variants in complex eye diseases. Cell genomics, 2(8), 2022.

19. F Alexander Wolf, Philipp Angerer, and Fabian J Theis. SCANPY: large-scale single-cell gene expression data analysis. Genome biology, 19(1):15, 2018.

20. Surag Nair, Mohamed Ameen, Laksshman Sundaram, Anusri Pampari, Jacob Schreiber, Akshay Balsubramani, Yu Xin Wang, David Burns, Helen M Blau, Ioannis Karakikes, et al. Deep learning the cis-regulatory code of chromatin dynamics during cellular reprogramming. bioRxiv, pages 2023–10, 2023.

21. Yong Zhang, Tao Liu, Clifford A Meyer, Jérôme Eeckhoute, David S Johnson, Bradley E Bernstein, Chad Nusbaum, Richard M Myers, Myles Brown, Wei Li, et al. Model-based analysis of ChIP-Seq (MACS). Genome biology, 9(9): R137, 2008.

22. Vikram Agarwal and David R Kelley. The genetic and biochemical determinants of mRNA degradation rates in mammals. Genome biology, 23(1):245, 2022.

23. Ziru Chen, Shijie Chen, Yuting Ning, Qianheng Zhang, Boshi Wang, Botao Yu, Yifei Li, Zeyi Liao, Chen Wei, Zitong Lu, et al. Scienceagentbench: Toward rigorous assessment of language agents for data-driven scientific discovery. arXiv preprint arXiv:2410.05080, 2024.

24. Siba Smarak Panigrahi, Jovana Videnović, and Maria Brbić. Heurekabench: A benchmarking framework for ai co-scientist. arXiv preprint arXiv:2601.01678, 2026.

25. Domen Jemec. GPTomics/bioSkills. https://github.com/GPTomics/bioSkills, Mar 30 2026. URL https://github.com/GPTomics/bioSkills.

26. Timothy Kassis, Clayton Young, Vinayak Agarwal, Jeremy Leipzig, Andrey Fedorov, Haoxuan “Orion” Li, urabbani, Cong, Jacob Luke, Salman Chishti layla, Alif Munim, Alper Yilmaz, Alyshia Ledlie Claude, Corin Wagen eamon-cervolve, Marek Wiewiórka, Yx Jiang, ashrafkahoush-ux, backtrue, Kuan, jiaodu1307, marovole, renato-umeton, and shao-shuai. K-Dense-AI/claude-scientific-skills. https://github.com/K-Dense-AI/claude-scientific-skills, Apr 3 2026. URL https://github.com/K-Dense-AI/claude-scientific-skills.

27. Marianne K DeGorter, Page C Goddard, Emre Karakoc, Soumya Kundu, Stephanie M Yan, Daniel Nachun, Nathan Abell, Matthew Aguirre, Tommy Carstensen, Ziwei Chen, et al. Transcriptomics and chromatin accessibility in multiple african population samples. bioRxiv, 2023.

28. 1000 Genomes Project Consortium et al. A global reference for human genetic variation. Nature, 526(7571):68, 2015.

29. Asa Thibodeau, Alper Eroglu, Christopher S McGinnis, Nathan Lawlor, Djamel Nehar-Belaid, Romy Kursawe, Radu Marches, Daniel N Conrad, George A Kuchel, Zev J Gartner, et al. AMULET: a novel read count-based method for effective multiplet detection from single nucleus ATAC-seq data. Genome biology, 22(1):252, 2021.

30. Nils Krietenstein, Sameer Abraham, Sergey V Venev, Nezar Abdennur, Johan Gibcus, Tsung-Han S Hsieh, Krishna Mohan Parsi, Liyan Yang, René Maehr, Leonid A Mirny, et al. Ultrastructural details of mammalian chromosome architecture. Molecular cell, 78(3):554–565, 2020.

31. Job Dekker, Andrew S Belmont, Mitchell Guttman, Victor O Leshyk, John T Lis, Stavros Lomvardas, Leonid A Mirny, Clodagh C O’shea, Peter J Park, Bing Ren, et al. The 4D nucleome project. Nature, 549(7671):219–226, 2017.

32. Peter Kerpedjiev, Nezar Abdennur, Fritz Lekschas, Chuck McCallum, Kasper Dinkla, Hendrik Strobelt, Jacob M Luber, Scott B Ouellette, Alaleh Azhir, Nikhil Kumar, et al. HiGlass: web-based visual exploration and analysis of genome interaction maps. Genome biology, 19(1):125, 2018.

